# An allometric scaling approach to estimate epiphytic bryophyte biomass in tropical montane cloud forests

**DOI:** 10.1101/2020.02.01.928515

**Authors:** Guan-Yu Lai, Hung-Chi Liu, Ariel J. Kuo, Cho-ying Huang

## Abstract

Epiphytic bryophytes (EB) are some of the most commonly found plant species in tropical montane cloud forests, and they play a disproportionate role in influencing the terrestrial hydrological and nutrient cycles. However, it is difficult to estimate the abundance of EB due to the nature of their “epiphytic” habitat. This study proposes an allometric scaling approach to measure EB biomass, implemented in 16,773 ha tropical montane cloud forests of northeastern Taiwan. A general allometry was developed to estimate EB biomass of 100 cm^2^ circular-shaped mats (n = 131) and their central depths. A point-intercept instrument was invented to measure the depths of EB along tree trunks (n = 210) below 3-m from the ground level (sampled stem surface area [SSA]) in twenty-one 30 × 30 m plots. Biomass of EB of each point measure was derived using the general allometry and was aggregated across each SSA, and its performance was evaluated. Total EB biomass of a tree was estimated by referring to an *in-situ* conversion model and was interpolated for all trees in the plots (n = 1451). Finally, we assessed EB biomass density at the plot scale and preliminarily estimated EB biomass of the study region. The general EB biomass-depth allometry showed that the depth of an EB mat was a salient variable for biomass estimation (R^2^ = 0.72, *p* < 0.001). The performance of upscaling from mats to SSA was satisfactory, which allowed us to further estimate mean (± standard deviation) EB biomass of the 21 plots (272 ± 104 kg ha^-1^) and to provide preliminary estimation of the total EB biomass of 4562 Mg for the study region. Since a significant relationship between tree size and EB abundance is commonly found, regional EB biomass may be mapped by integrating our method and three-dimensional airborne data.

## 1 INTRODUCTION

Bryophytes are rootless, non-vascular terrestrial plants such as mosses, liverworts and hornworts. Due to their primitive physiological characteristics, bryophytes are sensitive to the recent changes in climate such as increases in air temperatures (Aptroot & Van Herk 2007; Zotz & Bader 2009) and atmospheric carbon dioxide (Turetsky 2003), and decreases in precipitation (Gignac 2001). Epiphytic bryophytes (EB) are species that grow on the surface of a plant above the ground. They are some of the most representative lifeforms of tropical montane cloud forests (TMCF) (Barkman 1958; Smith 1982), which are ecosystems that experience frequent immersion of low altitude cloud (also known as “fog”, exchangeably used hereafter) with high humidity. Tropical montane cloud forests, as suggested by their name, are mostly distributed over mountainous regions. While covering only about 0.14% (∼30M ha) of the Earth’s terrestrial surface (Bruijnzeel, Mulligan, & Scatena 2011) and 2.5% of tropical forests of the world (Bubb *et al*. 2004), they are the major water sources for lowland environments. As a result, TMCFs play a disproportionately-large role in the functioning of a global terrestrial ecosystem relative to their limited distribution.

Epiphytic bryophytes may obtain necessary water and nutrients for growth by intercepting parallel fog water (Stadtmüller 1987; Holwerda *et al*. 2010; Scholl, Eugster, & Burkard 2011). In some regions, EB are keystone species for providing water and essential nutrients to maintain the health of TMCFs (Gradstein 2008; Zotz & Bader 2009) and may affect carbon storage of an entire ecosystem. They may also influence the global hydrological cycle by modifying precipitation and evaporation levels (Rhoades 1995; Chang, Lai, & Wu 2002; Porada, Van Stan, & Kleidon 2018). In the recent decades, land use and land cover change (Ray *et al*. 2006), and the prevailing global trend of elevated temperatures (Still, Foster, & Schneider 1999; Foster 2001) may alter regional climate in tropics, resulting in substantial ramifications on EB (Benzing 1998) and eventually TMCF. As “canaries in the coal mine” (Gignac 2001), spatiotemporal dynamics of EB may be effective indicators for monitoring the regional and global climate changes. One of the very first steps in this research field is to quantify the abundance of EB, which has been a very challenging task due to nature of their habitats and diverse morphologies (McCune & Lesica 1992).

Biomass is a major metric to assess the abundance of plants (Bonham 2013). For EB, biomass is also a key indirect parameter to assess the capacity of TMCFs to intercept fog (Zotz & Vollrath 2003). The abundance of EB in TMCFs may be affected by microclimatic (e.g., humidity, temperature, luminosity) and host structural (such as tree size, height and density) attributes (Peck, Hong, & McCune 1995; Freiberg & Freiberg 2000; Nöske *et al*. 2008; Chen, Liu, & Wang 2010). Field survey approaches such as destructively sampling with interpolation on the ground for low stature (Ah-Peng *et al*. 2017) or fallen (Chen, Liu, & Wang 2010) trees, and using a ladder, rope (Hsu, Horng, & Kuo 2002; Nakanishi *et al*. 2016), high tower or crane (McCune *et al*. 1997; McCune *et al*. 2000) to reach tall trees have been commonly implemented to measure EB biomass (see Table 1 a comprehensive summary). However, field EB measurements have been known to be quite challenging to carry out, which made regional quantification impractical (Moffett & Lowman 1995; Barker & Pinard 2001). In this paper, we proposed a simple and effective field allometric scaling method to estimate EB biomass for TMCF, which combines small-scale destructive field biomass collection, vertical point intercept sampling conducted by a newly-invented instrument, and up-scaling the biomass estimation with a previously established *in-situ* equation and data interpolation.

**TABLE 1.** Summary of the plot or the forest stand scale epiphytic bryophyte (EB) biomass density (kg ha^−1^) research reported in refereed literature. For the sake of quality, only peer-reviewed articles are listed. The table is organized based upon the data collection methods; “Climbing” includes the use of rope or ladder, and “Ground” indicates EB samples were reachable from the ground or removed from fallen logs. We note that studies that combined terrestrial bryophyte biomass or did not specify the collection of EB biomass only are not listed in this table. Annual precipitation (AP, mm y^-1^), mean annual temperature (MAT, °C) and elevation (m a.s.l.) of each site were directly obtained from its corresponding article. If the information was missing, it was then obtained from the internet. The ecosystems labelled as TMCF could be tropical montane cloud forest, or other similar forest ecosystems including tropical montane rain forest or tropical montane moist forest. The ones categorized as TCF are temperate conifer forests. To make the comparison legitimate, dead EB and humus mass was not included in the estimation. Studies only sampled part of EB biomass of trees such as a tree trunk (e.g., Kürschner & Parolly, 2004) are also not listed here.

| Method | Location | AP | MAT | Elevation | Ecosystem | Tree sample | EB biomass | Reference |
| --- | --- | --- | --- | --- | --- | --- | --- | --- |
| Climbing | La Soufrière, Guadeloupe | 1780 | 26.3 | 1330 | TMCF | Not available | 12336 | Coxson (1991) |
|  | Mascarene Archipelago, Madagascar | 8000 | 24 | 1350 | TMCF | Not available | 9020 | Ah-Peng *et al.* (2017) |
|  | Santa Rosa de Cabal, Colombia | 1250 | 5.5 | 3700 | TMCF | 1 | 6850 | Hofstede, Wolf and Benzing (1993) |
|  | Olympic Mountains, US | 4700 | 9.6 | 179 | TCF | 3 | 6527 | Nadkarni (1984a) |
|  | Cordillera de Talamanca, Costa Rica | 5193 | 16.8 | 1555 | TMCF | 15 | 6225 | Köhler *et al.* (2007) |
|  | Monteverde,<br>Costa Rica | 2591 | 18.6 | 1480 | TMCF | 25 | 4058 | Nadkarni *et al.*<br>(2004) |
|  | Cordillera de<br>Talamanca,<br>Costa Rica | 2812 | 10.9 | 2900 | TMCF | 6 | 1921 | Hölscher *et al.*<br>(2004) |
|  | Fushan, Taiwan | 3600 | 18.2 | 750 | TMCF | 18 | 1740 | Hsu, Horng and<br>Kuo (2002) |
|  | Monteverde,<br>Costa Rica | 2591 | 18.6 | 1700 | TMCF | 4 | 945 | Nadkarni (1984b) |
|  | Northeast China | 1450 | -0.8 | 875 | TCF | Not<br>available | 507 | Ye, Hao and Dai<br>(2004) |
|  | The Tilaran<br>Range, Costa<br>Rica | 5380 | 17.7 | 1325 | TMCF | 6 | 206 | Häger and<br>Dohrenbusch<br>(2011) |
| Harvesting | Monteverde,<br>Costa Rica | 2591 | 18.6 | 1480 | TMCF | 9 | 2087 | Nadkarni *et al.*<br>(2004) |
|  | Yunnan, China | 1931 | 11.3 | 2500 | TMCF | 77 | 1663 | Chen, Liu and<br>Wang (2010) |
|  | Cordillera<br>Oriental,<br>Colombia | 1850 | 6 | 3650 | Bamboo | Not<br>available | 1281 | Tol and Cleef<br>(1994) |
|  | Rwenzor<br>Mountains,<br>Uganda | 2000 | 8.5 | 3230 | TMCF | 1 | 1000 | Pentecost (1998) |
|  | Marafunga<br>Basin, New<br>Guinea | 3985 | 13 | 2625 | TMCF | 42 | 940 | Edwards and Grubb<br>(1977) |
|  | Zamora<br>Chinchipe,<br>Ecuador | 2080 | 15.5 | 2093 | TMCF | 63 | 604 | Werner *et al.*<br>(2012) |
|  | Central French Guiana | 2500 | 27 | 288 | TMCF | 15 | 452 | Gehrig-Downie *et al.* (2011) |
|  | Cascade Range, US | 2450 | 9.2 | 655 | TCF | 42 | 323 | McCune (1993) |
| Ground | Southern Thailand | 2000 | 28.5 | 804 | Tropical forests | 51 | 126 | Chantanaorrapint and Frahm (2011) |
|  | North Wales, UK | 2187 | 10.3 | 98 | TCF | 16 | 87 | Rieley, Richards and Bebbington (1979) |
| Scaling | Chilan mountain, Taiwan | 3500 | 12.7 | 1680 | TMCF | 210 | 272 | This study |

## 2 MATERIALS AND METHODS

### 2.1 Study site

The study was focused on 16,773 ha TMCF of Chilan Mountain (24°98’N, 120°97’E) in northeastern Taiwan (the spatial boundary defined by referring to Schulz *et al*., 2017). The precipitation in summer and winter consists of mostly orographic precipitation and tropical cyclones (regionally known as typhoons), and the northeastern monsoon, respectively. Annual precipitation and mean temperature of the site are 3,500 mm y^-1^ and 12.7°C, respectively. The mean (± standard deviation [SD]) elevation of the site is 1680 ± 343 m a.s.l., and mean slope (± SD) is 38.2° ± 13.4° ranging from 0° to 88.7°. The rugged terrain faces regular moist wind from the Pacific Ocean resulting in frequent occurrences of upslope fog approximately 300+ days of a year and 38% of the time (Lai *et al*. 2006). This humid bioclimate harbors a substantial amount of EB. There were 49 and 24 species observed in mature old-growth and regenerated forests, respectively, by a preliminary local inventory (Chang, Lai, & Wu 2002). The primary vegetation type of the TMCF is conifer forest, dominated by hinoki cypress (*Chamaecyparis obtusa* var. *formosana*) and Japanese cedar (*Cryptomeria japonica*). Bryophytes are the dominant epiphytic species of the region, occupying 93.5% of the total biomass (Deng 2006).

### 2.2 The patch scale EB biomass sampling and model development

The first step was to derive a general allometry for EB biomass, and six sites along the elevation gradient of 1200–1950 m a.s.l. were selected for sample collection (Figure S1). In the summer (May-October) of 2017, the center depth (e.g., from rhizoids to the top of a plant) of each EB species (n = 131; 113 liverworts, 17 mosses and 1 lichen) (for details of the species see the spreadsheet in Supplementary Information) within a randomly-selected 100 cm^2^ circular patch of a tree stem below 3 m above the ground was measured using a stainless steel ruler, and the sample was removed using a gardening shovel. Only a single species in the patch with the homogeneous depth was confirmed before the sample removal. The method has been applied previously by Rodríguez-Quiel, Mendieta-Leiva and Bader (2019). We note that one lichen sample was included in the model development due to the presence of a small portion of lichen among EB. The samples were stored in sealed linear low-density polyethylene bags to maintain moisture, then placed in an ice box and transported to a laboratory within eight hours after their removal from host trees. The samples were cleaned of dead organic matter, suspended soil and tree bark with tap water, dried in a 70°C biomass oven for at least 72 hours, and weighed using a three decimal place electronic balance (LIBROR EB-430H, Shimadzu, Japan). In this study, EB biomass was defined as the total sampled dry weight divided by the projected surface area of the sample (mg cm^-2^). The depth of EB was used as a unique trait for each independent sample to develop EB biomass allometric equations:

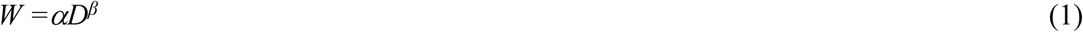

where *W* is the EB biomass (mg cm^−2^), *D* is the EB depth (cm), and *α* and *β* are the exponent components for the model. A power model was selected to fit the data by referring to previous studies (Niklas 1993; Niklas 2006) using R v. 3.5.0. (Stanford University; http://www.r-project.org/). Consecutive values ranging from 0.01 to 2.0 with an interval of 0.01 were selected for β with and without a fixed *α* value of 10 to derive an optimized model to fit the empirical data using generalized least squares. The method (generalized least squares) was specifically designed to minimize the effect of unequal variances, which were commonly observed in ecological data (Pinheiro & Bates 2006). Three variance covariate functions, the exponential of a variance covariate (varExp in R), power of a variance covariate (varPower) and constant plus power of a variance covariate (varConsPower), were used to modify regression of the fitted values and the residuals within the fitted model. The Akaike information criterion (AIC), the Bayesian information criterion (BIC) and log-likelihood were considered when facilitating model selection (Burnham & Anderson 2004). All statistical analyses were conducted using the “nlme” package in R (Pinheiro *et al*. 2019).

### 2.3 The tree scale EB biomass estimation

The main goal of this study was to implement a new field method for estimating EB biomass of TMCF at the regional scale. Once the allometric model (equation (1)) has been established, the next step was to estimate EB biomass of a tree, and we could then interpolate the estimate in the plot and regional scales. Twenty-one 30 × 30 m plots along the elevation gradient of 1260–1990 m a.s.l in Chilan Mountain of northeastern Taiwan were surveyed (Figure S1). Diameter at breast height (DBH) measured at 130 cm above the ground for each living tree with DBH ≥ 5 cm within 16 plots was recorded in July of 2016. The same approach was applied again to five more plots in January of 2019. During May-August of 2018 and January-February of 2019, we selected 10 trees (210 trees total) within each plot evenly distributed along the DBH gradient to interpolate EB biomass. Basal diameter (BD) of each sampled tree was also measured, and the relationship between basal area and DBH was investigated.

According to Johansson (1974) and Köhler *et al*. (2007), the majority of EB (in their case, 71–91%) were present at the lower part of a tree in TMCF, which may be utilized as a salient variable in estimating EB biomass of a tree. Therefore, a new field instrument was designed specifically for the estimation of EB biomass at the tree scale (Figure 1). From the ground to 300 cm of each sampled tree stem height, the EB depths (including the absence of EB with the depth of 0 cm) were recorded for every 30 cm vertical interval in several directions and were converted to biomass by referring to the allometry (equation (1)) and then averaged. The procedure was not vice versa due to the non-linearity of the allometry (a power model). We note that all trees in the plots were taller than 300 cm. The biomass of EB below 300 cm of a host tree was derived by taking the sampled stem surface area (SSA) into account. According to the visual inspection, the shape of the trunk from the ground to 130 cm was defined as a truncated cone and from 130 cm to 300 cm from the ground as a cylinder. Accordingly, the surface area (cm^2^) of the trunk below 3 m (SSA) was calculated by referring to equations (2) and (3):

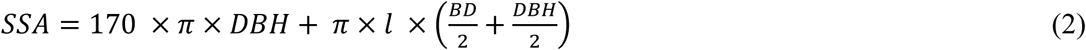

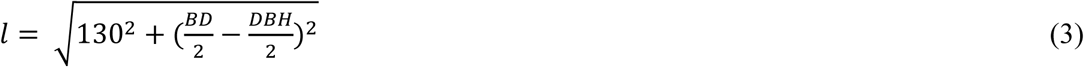

where SSA (cm^2^), *l* (cm), DBH (cm) and BD (cm) are sampled stem area, slant length of the cone, diameter at breast height and basal diameter, respectively. The sampled trees with DBH larger than 20 cm were recorded in eight directions (north, northeast, east, southeast, south, southwest, west and northwest) otherwise in just four major cardinal directions by referring to a compass. In August 2019, we stripped EB mats of SSA from 30 randomly selected and widely-distributed trees of different sizes to verify the estimation.

**FIGURE 1.**
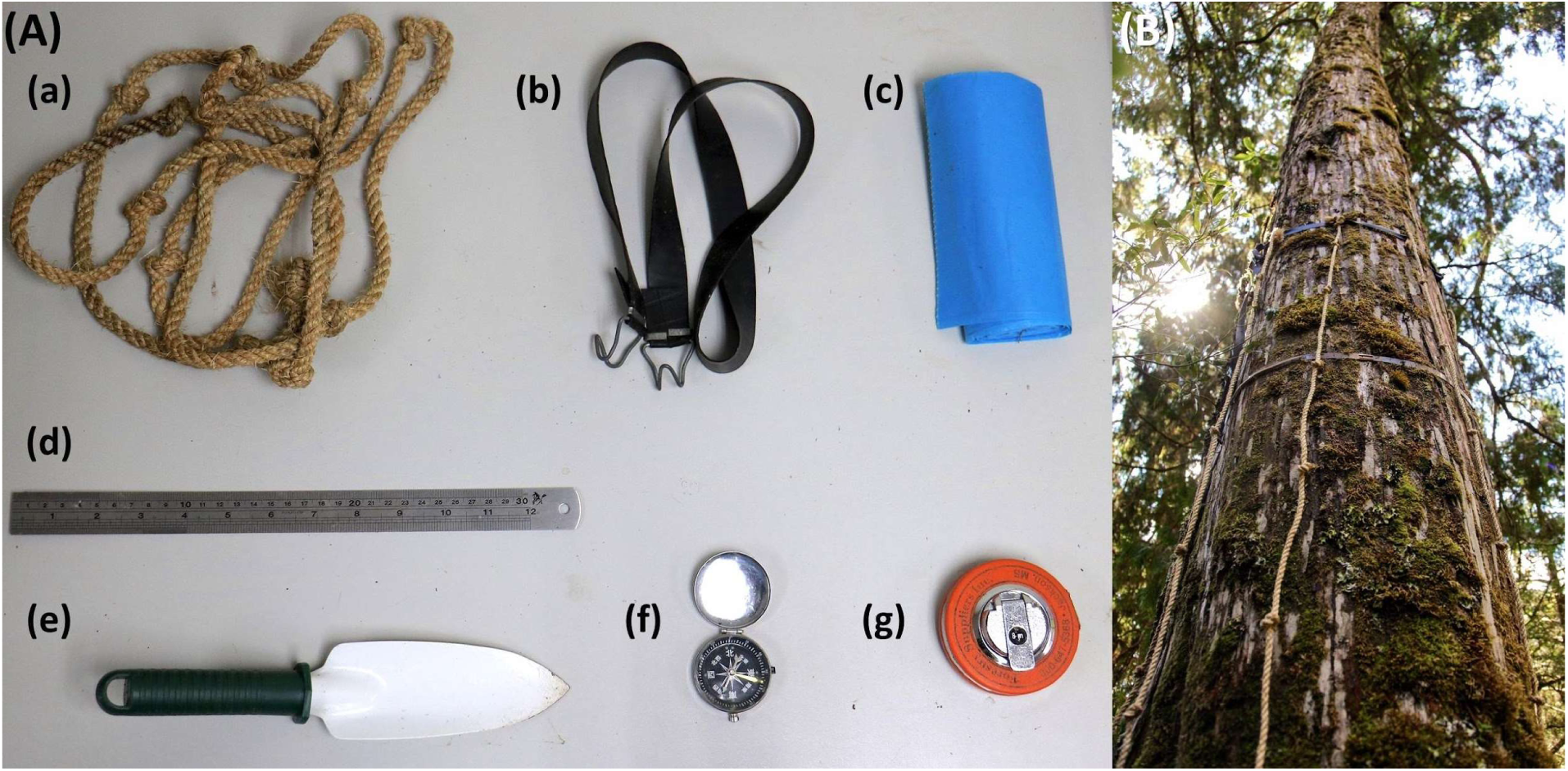
(A) The field instrument utilized in this study to estimate the biomass of epiphytic bryophytes (EB) in tropical montane cloud forests of northeastern Taiwan: (a) A 3-m rope with 30 cm long intervals marked by knots, (b) an adjustable rubber strip to fix ropes to a tree stem, (c) large, strong, and tear-resistant plastic bags to store EB from sampled stem surface area, (d) a stainless steel ruler to measure the heights of EB mats before removing samples with (e) a gardening shovel, (f) a compass to facilitate placing ropes in different orientations, (g) a fabric diameter tape to measure the sampled stem surface area. (B) A demonstration. The photograph was taken in Chilan Mountain by G. Lai in January 2019.

### 2.4 EB biomass up-scaling

The biomass of EB of 10 sampled tree was estimated by referring to equation (4):

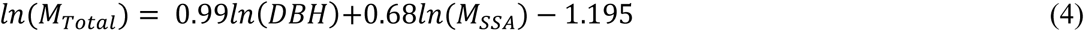

where *M*_*total*_ and *M*_*SSA*_ are EB biomass (kg) of total surface area and SSA of a tree, respectively, according to the *in-situ* destructive measurement by stripping EB from 10 harvested hinoki trees (R^2^ = 0.99, *p* < 0.001) (Deng 2006). Since the intercept of equation (4) is negative, resulting in negative values for small trees, a fixed ratio of 1.3 was then applied according to Deng (2006) for those trees. Sampled stem area of all trees (*S*_*total*_) in a plot was then estimated with the knowledge of DBH and DBH-BD of each tree (equations (2) and (3)), and EB biomass (*M*_*total*_, kg) (equation 5) and its density (kg ha^−1^) of a plot may be estimated by referring to equation (5) with the knowledge of EB biomass (*M*_*sampled*_) on SSA (*SSA*_*sampled*_) of 10 sampled trees.

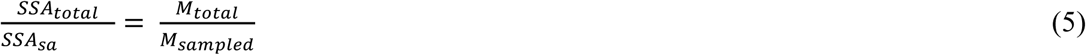

Literature search was conducted in Google Scholar (https://scholar.google.com/) with the keywords “epiphytic bryophyte” and “biomass” for a general comparison of EB biomass density (with basic bioclimatic information). We note that for the sake of quality control, non-refereed articles such as graduate theses and conference proceedings were excluded. Finally, since the stand characteristics of selected plots were quite representative of the region by referring to Wang and Huang (2012), Hu and Huang (2019) and several local inventory data, EB biomass of TMCF in Chilan Mountain may be estimated after taking the areal size of the region (16,773 ha) into account.

## 3 RESULTS

### 3.1 Epiphytic bryophytes biomass allometry

In this study, we collected 100 cm^2^ circular-shaped EB samples (n = 131) from six forest stands in Chilan Mountain along an elevation gradient. The mean (± SD [minimum–maximum]) sampled EB depth and biomass were 4.5 ± 2.9 cm (0.3–13.7 cm) and 36.0 ± 20.3 (6.2–99.3) mg cm^−2^, respectively. Significant positive correlations (*p* < 0.005) were found among EB depth and biomass with different regression models (Table 2). Performance of the allometric equation of the power of variance covariate function (R^2^ = 0.72, *p* < 0.0001) with smaller AIC and BIC and greater log likelihood was superior to other models, and the model was selected for further analyses (Figure 2).

**TABLE 2.** Model performance comparison of allometric equations (*W* = *αD*^*β*^, equation (1)) by referring to values of the Akaike Information Criterion (AIC), the Bayesian Information Criterion (BIC) and log likelihood. We note that all models are significant with *p* < 0.001.

| Model | $\alpha$ | $\beta$ | $R^2$ | AIC | BIC | Log likelihood |
| --- | --- | --- | --- | --- | --- | --- |
| Nonlinear squared regression | 12.62 | 0.72 | 0.72 | 395.12 | 403.75 | -194.56 |
| Nonlinear squared regression* | 10.00 | 0.84 | 0.70 | 400.34 | 406.09 | -198.17 |
| Power of a variance covariate | 11.96 | 0.75 | 0.72 | 379.92 | 391.43 | -185.96 |
| Exponential of a variance covariate | 11.77 | 0.77 | 0.72 | 380.12 | 391.62 | -186.06 |
| Constant power of a variance covariate | 11.78 | 0.76 | 0.70 | 380.91 | 395.28 | -185.45 |

\*Nonlinear squared regression with the fixed $\alpha$ of 10.00

**FIGURE 2.**
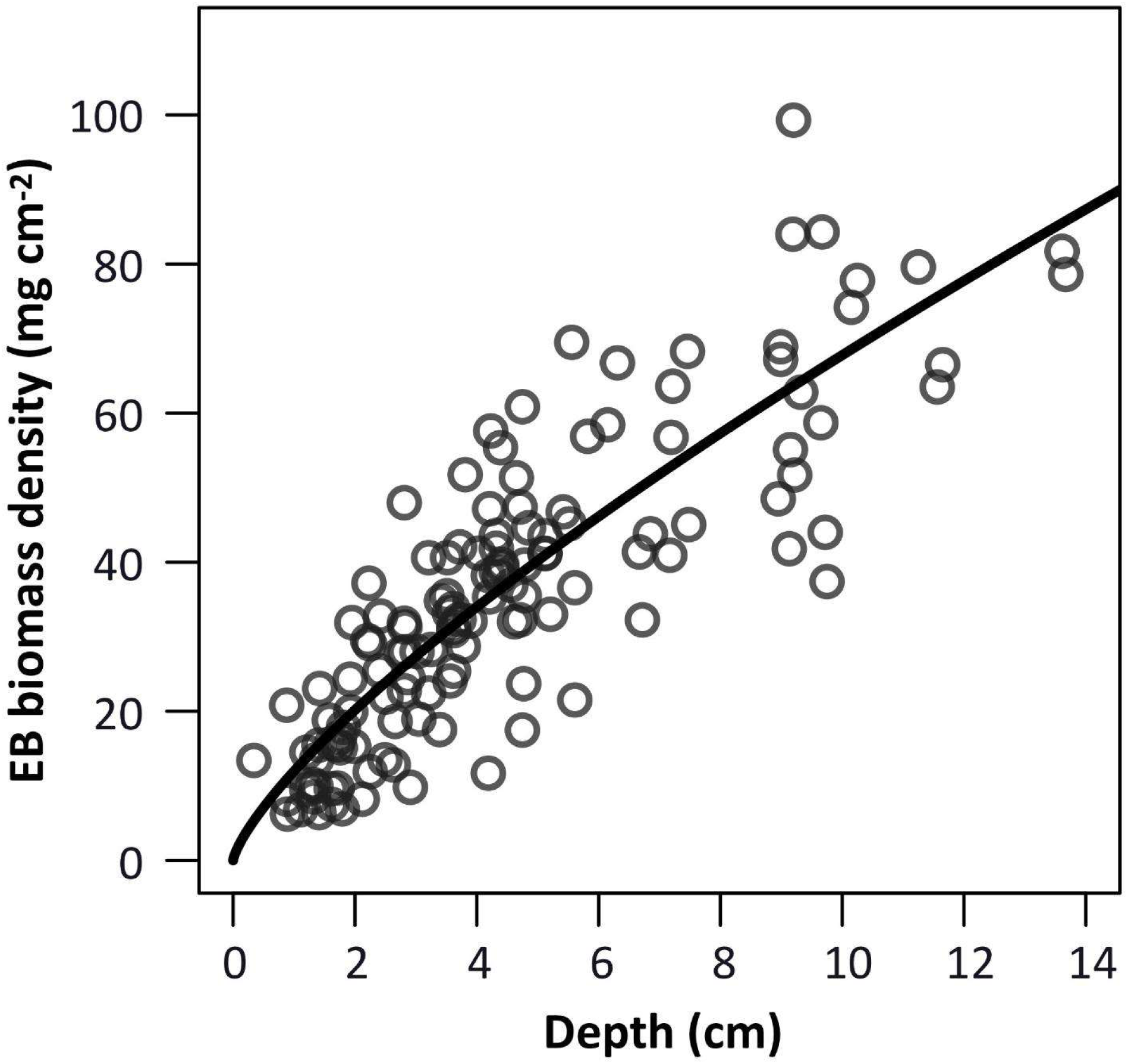
The best empirical general depth-biomass allometric model of epiphytic bryophytes (EB). The model was a power of variance covariate function (R^2^ = 0.72, AIC = 380, *p* < 0.001, n = 131), and the performance was superior to other models (Table 2) with coefficient and exponent of 11.96 and 0.75, respectively.

### 3.2 The tree-scale EB biomass estimation

Ten trees evenly distributed along the DBH gradient of each plot (total 210 trees) were selected to investigate the relationship between DBH and BD of EB-hosted trees. The mean (± SD [minimum–maximum]) DBH and BD of sampled trees were 33.5 ± 27.8 (7.6–128.7) cm and 49.5 ± 34.5 (9.9–186.2) cm, respectively. High correlation (R^2^ = 0.94, *p* < 0.0001) was found between DBH and BD (Figure S2). With this information, we computed SSA in the plots by referring to equations (2) and (3). The statistics (mean ± SD [minimum–maximum]) of SSA was 3.5 ± 2.8 (0.81–13.5) m^2^. Mean (± SD [minimum–maximum]) EB depth of the 210 sampled trees was 1.1 ± 0.6 (0.1–3.1) cm, and the data was injected into the allometry (Figure 2) to yield EB biomass (mean ± SD [minimum–maximum]) of 10.2 ± 5.2 (0.7–26.1) mg cm^−2^ (or 402.2 ± 478.9 [8.3–2856.6] g) on SSA. We note that there was a significant positive curvilinear relationship (*p* < 0.001) between DBH of the sampled tree and EB biomass on SSA (Figure 3).

**FIGURE 3.**
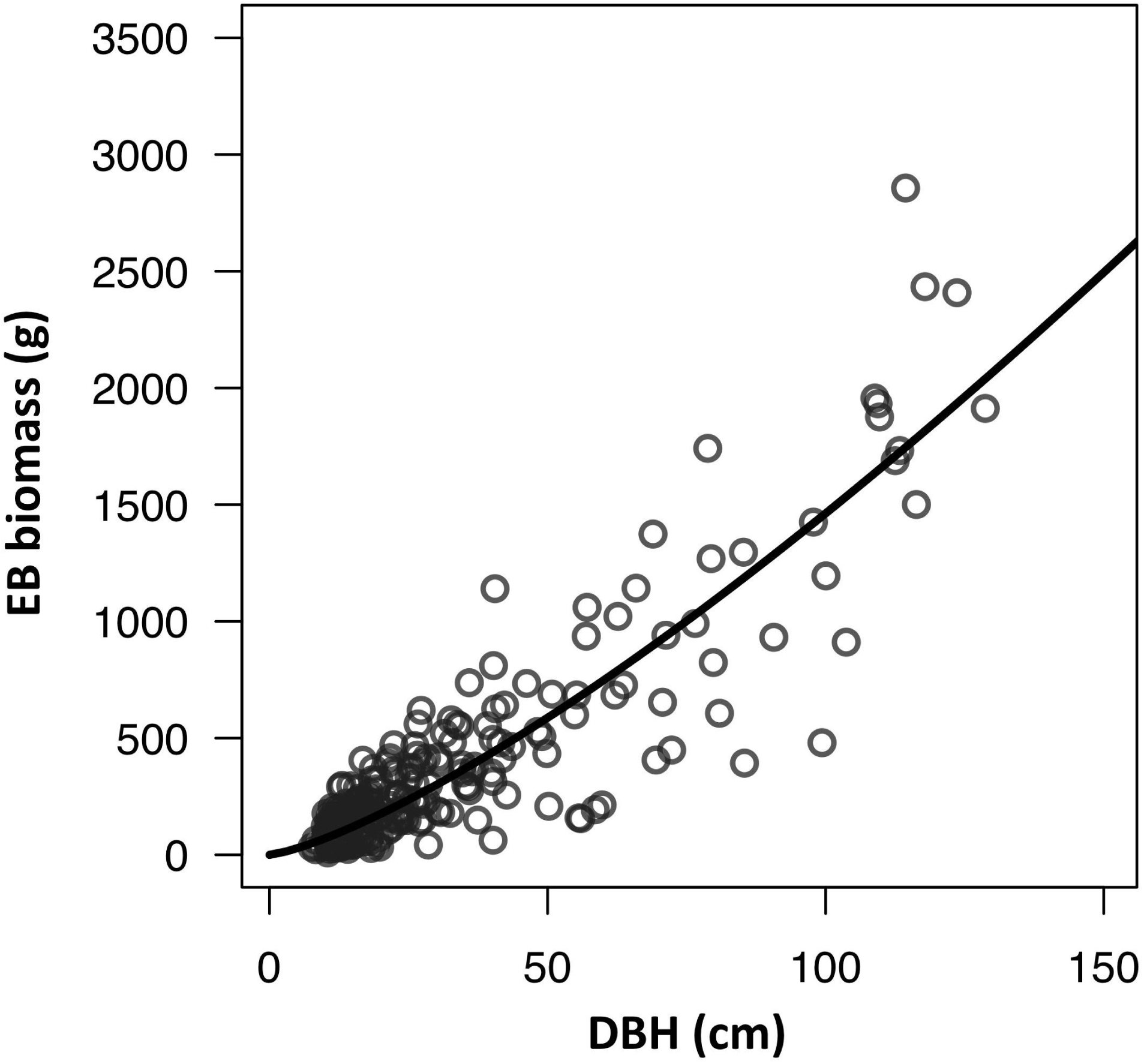
The relationship between diameter at breast height (DBH) and epiphytic bryophyte (EB) biomass of sampled stem surface area based upon 10 sampled trees of different DBH sizes on the 21 field plots (n = 210, Figure S1): EB biomass = 3.40DBH^1.32^ (R^2^= 0.86, *p* < 0.001).

Biomass of epiphytic bryophytes on 30 randomly selected trees with mean (± SD, minimum–maximum) DBH of 26.2 ± 21.5 (5.7–93.0) cm was destructively collected to verify the proposed approach of upscaling the patch scale estimation (Figure 2) to SSA. Overall, the performance was satisfactory (Figure 4) and all samples but one outlier (R^2^ = 0.82 and 0.95 without the outlier, *p* < 0.0001 for both model) were close to the 1:1 line (slope = 0.93 and 0.95 without the outlier, *p* > 0.8 for the intercepts of both models) with the mean absolute difference of 77.3 g (35.2% of the mean estimate) or 56.3 g (25.2% of the mean estimate) without the outlier. The outlier may be possibly due to rotten and soften tree barks underneath the EB mats (observed during the sample cleaning), and the depth of tree bark may have been included in the EB depth measurement, resulting in pronounced over-estimation. By applying the *in-situ* conversion function (equation (4)), the EB biomass (mean ± SD [minimum–maximum]) for each sampled tree within the plots was estimated (818.3 ± 1335.1 [12.9–7279.1]) g (n = 210).

**FIGURE 4.**
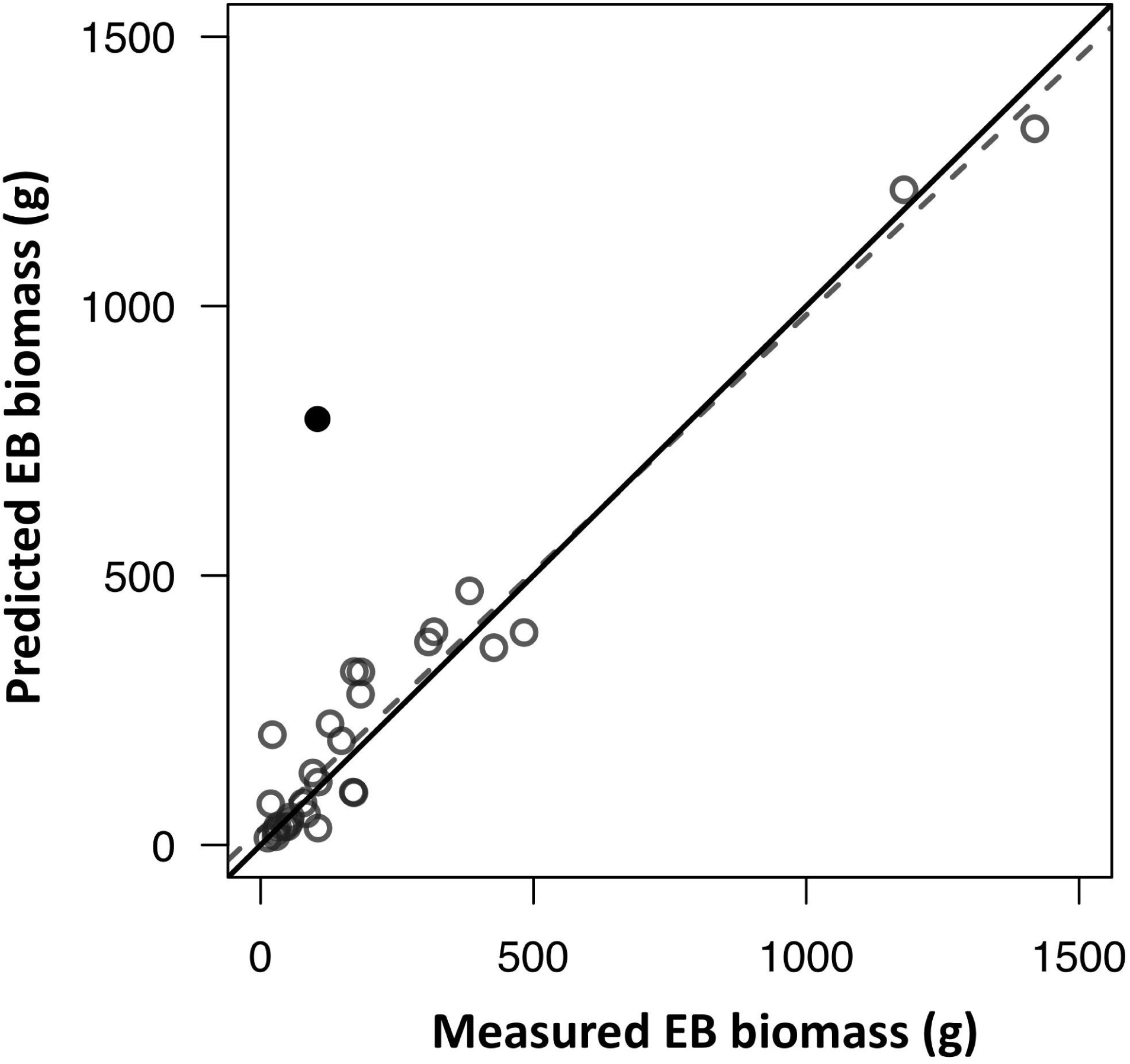
The comparison of model-predicted epiphytic bryophyte (EB) biomass and field collected EB biomass. The black solid dot indicates an apparent outlier in which EB inhabited on decomposed tree bark.

### 3.3 The plot and regional scales EB biomass estimation

Mean (± SD [minimum–maximum]) DBH of the trees (n = 1451) within twenty-one plots was 20.3 ± 17.5 cm (5.0–176.0 cm) (detailed plot-scale statistics of forest stands see Table S1). The EB biomass (and biomass density) for each plot can be interpolated by referring to the EB biomass of 10 sampled trees within each plot with the mean ± SD (minimum–maximum) of 24.5 ± 9.4 (8.8-39.0) kg (or 272.0 ± 104.0 [97.9–433.3] kg ha^−1^). Twenty-one refereed papers were found, and 86% (18/21) of the studies reported higher EB biomass density values than our mean plot/stand scale estimation (Table 1). Finally, with the knowledge of the plot-scale mean EB biomass density, we provided the preliminary estimation of the total EB biomass of 4562 Mg for the 16,773 ha TMCF of Chilan Mountain.

## 4 DISCUSSION

Epiphytic bryophytes are some of the most quintessential species characterizing mid-altitude tropical montane cloud forests (Bruijnzeel, Scatena, & Hamilton 2011) and play a pivotal role in influencing the global hydrological cycle (Porada, Van Stan, & Kleidon 2018). Due to the diverse morphology of the species and their “epiphytic” habitat, it is difficult to quantify the abundance of EB. In this study, we propose a novel field protocol for regional EB biomass estimation. Our discussion will mainly focus on (1) EB depth-biomass allometry, (2) scaling of EB biomass from the patch to the regional scale, and (3) limitation and future directions.

### 4.1 The patch scale EB depth-biomass allometry

In this study, *in-situ* general allometric equations were developed to estimate the biomass of a 100 cm^2^ circular patch of EB using the central depth of the sample (Figure 2). The performance was satisfactory, even though the morphology of EB is much more diverse than most vascular plants. Plant allometry focuses on relationships between plant body size and biomass, production, population density or other abundance related dependent variables (Enquist, Brown, & West 1998; Enquist *et al*. 1999). Stanton and Reeb (2016) suggested that some characteristics of bryophytes may be allometrically scaled like vascular plants, which was verified in this study. The mean exponent of the five selected power models was 0.75 (3/4) (Table 2), which agrees with the 3/4 power law (Kleiber 1947) and is similar to the constant scaling exponents over a wide range of vascular plant size, often with quarter-powers in metabolic scaling theory using biomass as an independent variable (West, Brown, & Enquist 1997; West, Brown, & Enquist 1999). However, epiphytic bryophytes are non-vascular plants composed of a simple stem, which has a limited role in transporting moisture and nutrients through conducting tissues and does not follow the vascular transport system as a self-similar, fractal-like branching network (Ligrone, Duckett, & Renzaglia 2000). Two major branching forms of bryophytes are sympodial with connected modules of equal level and monopodial (Stanton & Reeb 2016). For most vascular plants, the branching bifurcation is two (Enquist *et al*. 2007), and the height is 1/4 exponent of mass (West, Brown, & Enquist 1999). It was different to our empirical observation, although the sampling unit was a mat but not an individual. This could verify that the basic assumption of an organism’s self-similar branching network plays a major role in governing the allometric relationship.

### 4.2 Up-scaling of EB biomass

A point-intercept field instrument was invented in this study to facilitate sampling EB height data along a tree stem, which were then used as an independent variable to estimate EB biomass (Figure 2) and EB biomass of SSA, and later extrapolate to the tree scale using an *in-situ* conversion equation (Equation (4)). The distribution of EB biomass on a tree could be very sensitive to the ambient environment (McCune 1993; Sillett & Antoine 2004). Therefore, we measured the depth of EB in four and eight directions for small (DBH ≤ 20 cm) and large (DBH > 20 cm) trees, respectively, which may reduce microclimate-induced biases. The method was efficient, taking about 15 minutes for the four-direction measurement and double that amount of time for the eight-direction measurement. This may permit rapid sampling to obtain a large sample size (Table 1). With proper sampling design and data inter/extrapolation, we may be able to estimate EB biomass in a large region. Mean biomass density of EB estimated in this study was similar to the one conducted in the same region (230 kg ha^-1^) but within a much smaller spatial extent using a destructive tree harvesting approach (Deng 2006). Our mean plot (forest stand) scale estimation of EB biomass density falls within the lower half of the EB biomass density global synthesis data (Table 1). It is challenging to make a fair comparison since those previous studies were conducted using different data collection methods over a wide range of spatial extents. However, in terms of efficiency, the proposed new approach is indeed superior to other sampling methods implementing for the sampling of 210 EB host trees in this study.

This point-intercept approach should also be applicable for the estimation of ground bryophyte biomass, and facilitates the estimation of overall abundance of bryophytes in an ecosystem. This is a pivotal but rarely available parameter, and has a major impact on regulating the terrestrial hydrological cycles (Porada, Van Stan, & Kleidon 2018). This study focused on the height of a tree below 3 m from the ground, where the majority of EB are present (Trynoski & Glime 1982) (Figure 1B). The sampled stem area may be further extended with aids of a foldable ladder.

### 4.3 Limitation and future directions

One potential research limit is that the tree scale EB biomass estimation, which was extrapolated from the estimation on SSA (equation 4), could not be validated with empirical data. The task is rather difficult and may be impractical for the study region. It requires tree climbing or destructive tree harvesting to strip EB of an entire tree. However, the support of tree climbing was not available during the time of conducting this study, and it could be risky to climb a small-size tree without reliable support for a climber’s body weight. Logging for both natural and plantation forests has been completely forbidden in Taiwan since 1991. Therefore, the latter option may not be possible due to the local regulation. In the future, we might be able to take the advantage of tropical cyclone-induced fallen logs and harvest EB biomass at the ground level, since the island is located in a typhoon-prone region (Chi *et al*. 2015). However, this sampling approach could be biased since the probability of the strong wind induced tree falling may be associated with topography (Mitchell 2013), which also plays a pivotal role in governing the abundance of EB (Werner *et al*. 2012).

It is extremely challenging to non-destructively measure EB biomass, and a new field approach was developed in this study to tackle this task. This is crucial because the age of EB on a tree could be almost as old as the age of the host tree (Kimmerer 2003), and it may require many years of recovery after the removal of samples (Fenton, Frego, & Sims 2003). It may be useful to further generalize the EB allometry (see the supplementary spreadsheet data) to make it applicable for other settings. According to this study (Figure 3) and some previous literature (Hsu, Horng, & Kuo 2002; Köhler *et al*. 2007; Chen, Liu, & Wang 2010), we found that there may be a significant relationship between the tree size and the abundance of EB. With the availability of a three-dimensional tree size spatial layer at the regional scale derived from high spatial resolution airborne lidar (light detection and ranging) or aerial photographic point cloud data (Chung *et al*. 2019; Kellner *et al*. 2019), we may be able to map EB biomass over a vast region.

## ACKNOWLEDGEMENTS

We appreciate Jun Zhang and Hong-You Lin for providing field assistance. This study was sponsored by the Ministry of Science and Technology of Taiwan (MOST 106-2633-M-002-002-), National Taiwan University EcoNTU project (106R104516), and the NTU Research Center for Future Earth from the Featured Areas Research Center Program within the framework of the Higher Education Sprout Project by the Ministry of Education in Taiwan.

## AUTHORS’ CONTRIBUTIONS

GYL and CyH conceived the idea and developed the method for this research and led the writing of the manuscript; GYL, AJK and HCL analyzed the data. All authors collected field data and contributed critically to the drafts and gave final approval for publication.

## DATA AVAILABILITY

Data used to derive epiphytic bryophyte allometry can be found in Supplementary Information.

## Notes

http://homepage.ntu.edu.tw/~choying/PJ_SI2.xlsx

