## Supplementary Information for "An allometric scaling approach to estimate epiphytic bryophyte biomass in tropical montane cloud forests"

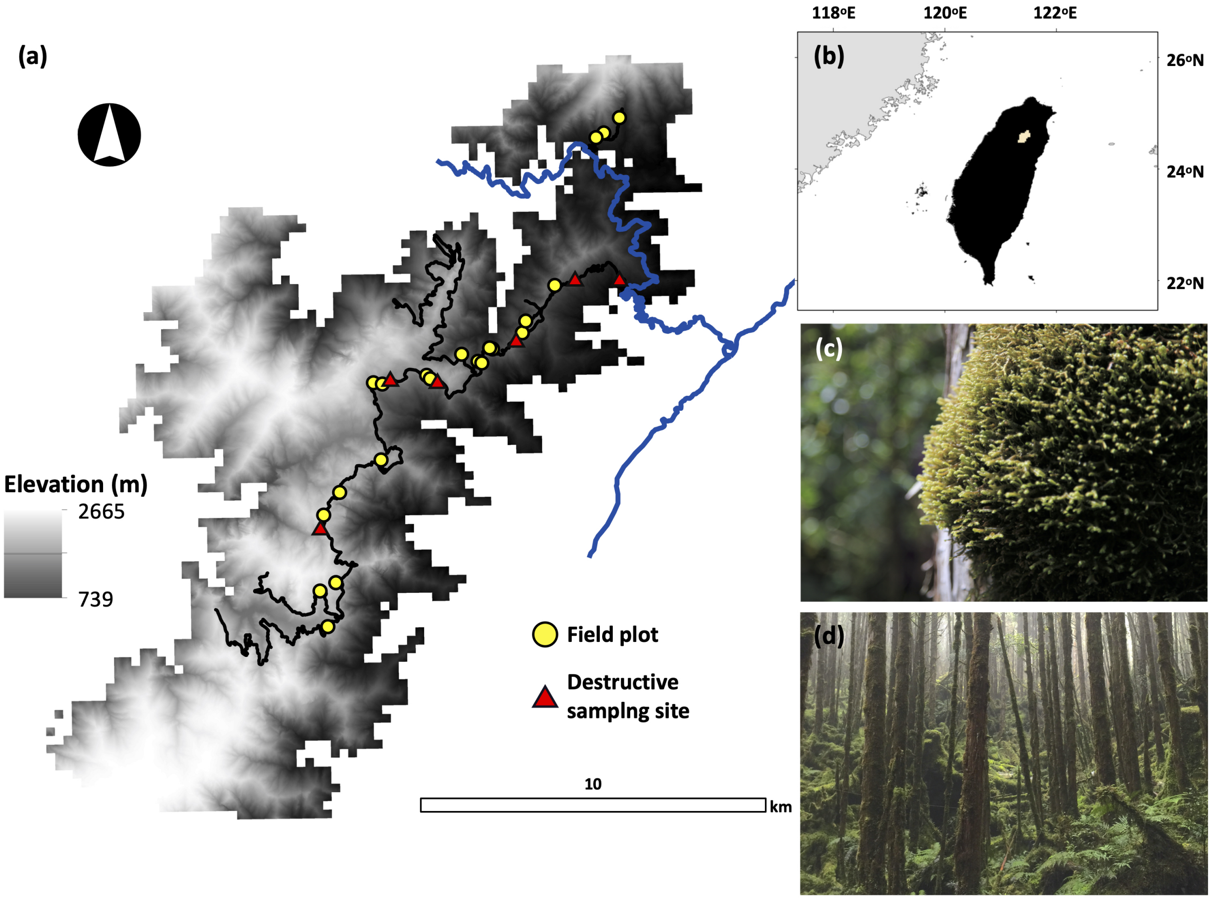


**FIGURE S1** (a) The study area (24°98'N, 120°97'E), a tropical montane cloud forest in Chilan Mountain, located in (b) northeast Taiwan (the yellow polygon). The blue and black colored lines are the state and unpaved forest routes, respectively. The gray monochromatic colored background is the 30 m ASTER (the Advanced Spaceborne Thermal Emission and Reflection Radiometer) Global Digital Elevation Model data. Epiphytic bryophyte (EB) mat samples (n = 131, Figure 2) were destructively collected from six forest stands (red triangles) to develop depth-biomass allometric models to estimate EB biomass in twenty-one 30 × 30 m plots (yellow dots) along the elevation gradient. (c) *Bazzania* spp. is one of the dominant EB species found in (d) this tropical montane cloud forest. The photographs were taken by GY Lai in summer 2017.

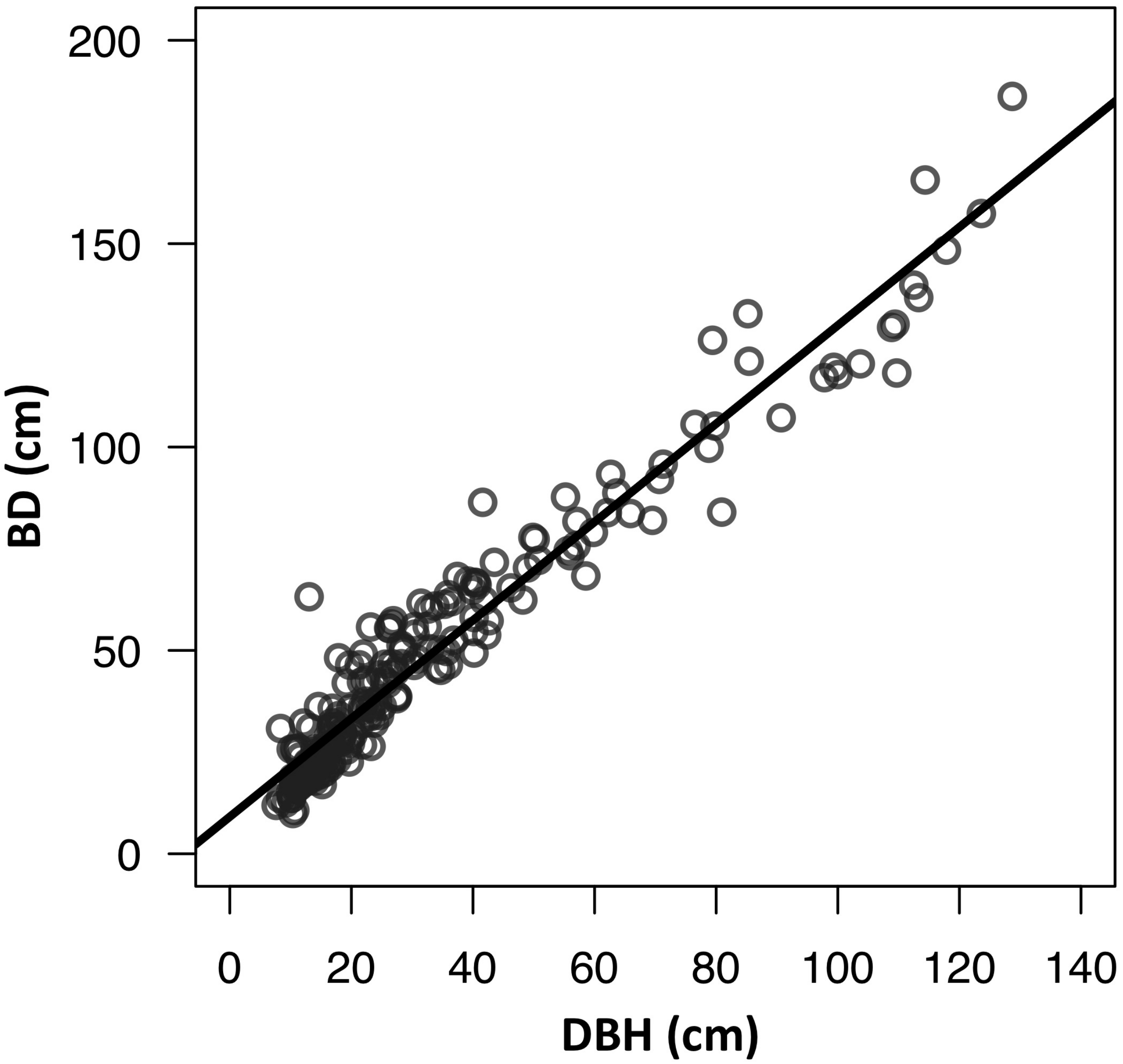


**FIGURE S2** The relationship between diameter at breast height (DBH) and basal diameter (BD) of 210 sampled trees within twenty-one field plots (BD = 9.1 + 1.2DBH, $R^2 = 0.94$, $p < 0.001$).

**TABLE S1** Statistics (mean, standard deviation [SD], minimum [min], maximum [max]) of twenty-one 30 × 30 m forest stand characteristics (count, density, diameter at breast height [DBH] and sampled stem surface area below 3 m [SSA]).

| | Mean (SD) | Min–Max |
|---|---|---|
| Count | 69.1 (46.3) | 9.0–208.0 |
| Density (# $ha^{-1}$) | 767.7 (514.9) | 100.0–2311.0 |
| DBH (cm) | 28.6 (18.8) | 5.0–175.0 |
| SSA ($m^2$) | 143.6 (58.6) | 62.6–284.1 |
